# TGS-GapCloser: fast and accurately passing through the Bermuda in large genome using error-prone third-generation long reads

**DOI:** 10.1101/831248

**Authors:** Mengyang Xu, Lidong Guo, Shengqiang Gu, Ou Wang, Rui Zhang, Guangyi Fan, Xun Xu, Li Deng, Xin Liu

## Abstract

The completeness and accuracy of genome assemblies determine the quality of subsequent bioinformatics analysis. Despite benefiting from the medium/long-range information of third-generation sequencing techniques, current gap-closing tools to enhance assemblies suffer multi-alignments and high error rates, resulting in huge time and money costs.

We developed a software tool, TGS-GapCloser that uses the low depth (>=10X) single molecule sequencing long reads without any error correction to close gaps. The algorithm distinguishes gap regions from the alignments of long reads against original scaffolds, corrects only the candidate regions, and assigns the best sequences to each gap. We demonstrate that TGS-GapCloser improves the contig N50 value of draft assembly by 25-fold on average, updating above 90% gaps with 93.96% positive predictive value. Despite of high error rate of raw long reads, improved assemblies archive Q50 (99.999%) single-base accuracy with only 11.8% decrement to inputs. Besides it could complete more gaps, and is also ∼29-fold faster than mainstream gap-closing tools. BUSCO analysis revealed that 3.4%-13.1% more expected genes were complete. TGS-GapCloser also shows its power to fill gaps for ultra large genome assembly of ginkgo (∼12Gb) with 71.6% of gaps closed. The validation of inserted or merged gap sequences was conducted with NGS reads and reference genomes, respectively. The updated genome assemblies may promote the gene annotation, structure variant calling and thus improving the downstream analysis of ontogeny, phylogeny, and evolution.

## 1 Introduction

### 1.1 Gap closure perfects current genome assemblies

The development of genome sequencing techniques has reduced the cost and improved the throughput at a speed beyond Moore’s Law over the last decade[1]. The genetic sequence databases have been drastically enriched, and progressively increasing focuses move from small bacterial and fungal genomes to large eukaryotes. The applications of state-of-the-art techniques, for instance, third-generation long reads (TGS)[2, 3], synthetic long reads (SLR)[4-8], Hi-C[9], and BioNano physical map[10], provide extra information on different length scales, resulting in the enhanced genome assemblies. However, all the complete assemblies are imperfect, even for human and model organisms, which contain unknown nucleic acids (represented by Ns), as the Bermuda Triangles areas in the scaffold island chains. The repetitiveness and polymorphism of the genomes, the limitation of sequencing techniques, and the trade-off of algorithms may lead to the Bermuda areas. Gap closure or gap filling can discover the sequences and extend contigs to entirely or partially missing gene-encoding area. Therefore, there is a need to develop tools to close gaps in *de novo* genome assemblies, and obtain more complete and accurate genomes, especially for large eukaryotic genomes with high complexity.

### 1.2 Problems in Sangers/ NGS gap-closing tools

The first effort to finish the gaps in draft genome assemblies was made using Fosmids, BACs libraries and Sanger reads for a large range of 1 to 100 kb[11]. But the manual or semi-automated processes limit the applications in consideration of huge costs. The next-generation sequencing (NGS) technologies along with paired-end and mate-pair information of multiple insert sizes have overcome the financial problem, and several benchmarking tools have been designed to reach into gap regions [12-16], sharing similar kmer-extension or local reassembly algorithms, but suffer the same problems of CPU hour and memory consuming for large genomes. Besides, those strategies cannot span the repetitive DNA fragments such as tandem repeats, fail to handle with overlapped neighboring contigs in most cases and cause more misassemblies due to the short read/kmer length.

### 1.3 Problems in current TGS assemblies and TGS gap-closing tools

The single molecule TGS technologies, including Pacific Biosciences (Pacbio) and Oxford Nanopore Techniques (ONT) have the potential to solve these limitations as their reads (∼10kb) are typically longer than most DNA repeats[17]. Although *de novo* genome assembly using long reads may allow incremental improvements, the higher expense and lower accuracy relative to NGS platforms prevent the popularization. Common TGS sequencing errors, insertions or deletions, may cause frameshifts in gene-coding regions, disrupting the gene annotations. There have been several hybrid assemblers designed to combine advantages of both sequencing platforms since the TGS commercial platforms were released. Main principles include indiscriminately constructing final assembly graph by mixing NGS contigs and TGS long reads based on OLC or string graph algorithm[18], or scaffolding short contigs generated by NGS dependent on their alignments on long reads[19, 20] to utilize the medium-range information, but ignore the possible combination with long-range information provided by other techniques. Gap-closing algorithms, however, only upgrade the missing regions, reserving the majority of the existing assembly to considerably reduce the computing complexity and cost. PBJelly[21] was the first software to use PacBio dataset to close gaps through locally assembling the mapped long reads in gap regions. The number of gaps could also be efficiently reduced by FGAP[22], which aligned long reads to the gaps using BLAST algorithm[23]. More tools modified the algorithm and extended for different purposes[24-28]. However, most tools mentioned above share the same crucial shortcoming: they only accept pre-error-corrected long reads or alternative assembled contigs. It hampers the application of long-range information because the error correction using TGS data themselves needs sufficient coverage of expensive long reads, usually splitting them into short fragments and losing valuable length information, while that using NGS data requires huge memory consumption, not readily usable for large genomes.

### 1.4 Key points in the design of TGS gap-closing tools

Three key points are need to be considered to develop a TGS gap-closing algorithm. First, use TGS data as few as possible. Although the price of TGS has been decreasing[29], the efficiency is still the first priority, especially for those small labs or small projects. Thus, local reassembly or pre-error correction by overlapping long reads is not preferable. Another important factor is the accuracy in choices of long reads to fill the gaps. It has been demonstrated that the number of assembly errors caused by most gap-closing tools is higher than that of *de novo* assembled contigs [25]. High error rate and the existence of repeats may increase the probability of large misassembly events. Last but not least, the filled gaps should not diminish the single-base level accuracy, which determines the quality of downstream analysis. There is still a need of error correction or polish for inserted sequences. Note that the most recent Pacbio improved its base-calling accuracy to 99.8%[30], which may directly simplify the problem, but drastically sacrifice the throughout and read length. In this work, we described a software tool, named TGS-GapCloser, that uses error-prone long reads at low coverage to efficiently and accurately close gaps within a reasonable time. We applied it to three draft genome assemblies of human using ONT or Pacbio long reads[31, 32], improving the contig NG50 5.5 to 44.9-fold and NGA50 5.1 to 30.7-fold, finishing above 90% gaps with 93.96% positive predictive value (PPV) and 65.97% sensitivity on average. 71.6% gaps in the ultra large genome assembly of ginkgo were also closed using 11.9 coverage of corrected Pacbio long reads, increasing the contig N50 from 48kb to 365kb.

## 2 Methods

### 2.1 Genome assemblies and TGS datasets

Three datasets of two large genomes were used to examine the gap-closing results by TGS-GapCloser. We sequenced *Homo sapiens* (NA12878) using MGIEasy single tube-Long Fragment Reads (stLFR) Library Prep Kit on BGISEQ-500 platform with the data size of 660 Gb, and reads mapped to the Chromosome 19 (Chr19) were also extracted for further analysis. NGS short reads were assembled using *de Bruijn* graph-based assemblers, MaSuRCA[20] and Mercedes (inhouse tool) to obtain short but highly accurate contigs for each dataset, and the SLR long-range and paired-end information provided by stLFR technique was exploited to do further scaffolding by SLR-superscaffolder[31]. Supernova[33] was originally designed to assemble 10X Genomics data, but could be applied to stLFR format reads to obtain draft scaffolds. To test the generalization of TGS-GapCloser, we utilized both Pacbio RSII (SRR3197748) downloaded from GIAB and ONT MinION (rel3)[32] long reads to close the gaps.

The input genome assembly of *Ginkgo biloba* female (estimated 12 Gb) was obtained from [34], which assembled using SOAPdenovo2[12] and updated using Hi-C data. The Pacbio reads for ginkgo were sequenced by Pacbio Sequel, with chemistry of Sequel Sequencing Kit 3.0 Bundle (4 rxn). The total data amount was 256 Gb with the average read length of 38,623 bp. Error-correction by Canu[35] reduced data to 126 Gb, with the average read length of 10,722 bp.

### 2.2 Algorithm of TGS-GapCloser

TGS-GapCloser accepts any kind of TGS long reads or other pre-assembled contigs to automatically fill gaps in any kind of draft assembles in the following four steps shown in Figure 1: (i) determination of gap regions in the draft assembly; (ii) acquisition of candidates from the alignments of long reads against gaps; (iii) base-level error correction of alternative sub-long reads; and (iv) gap closure with the error-corrected candidates with the highest score for each gap or linkage of the neighboring contigs with overlap.

**Figure 1.**
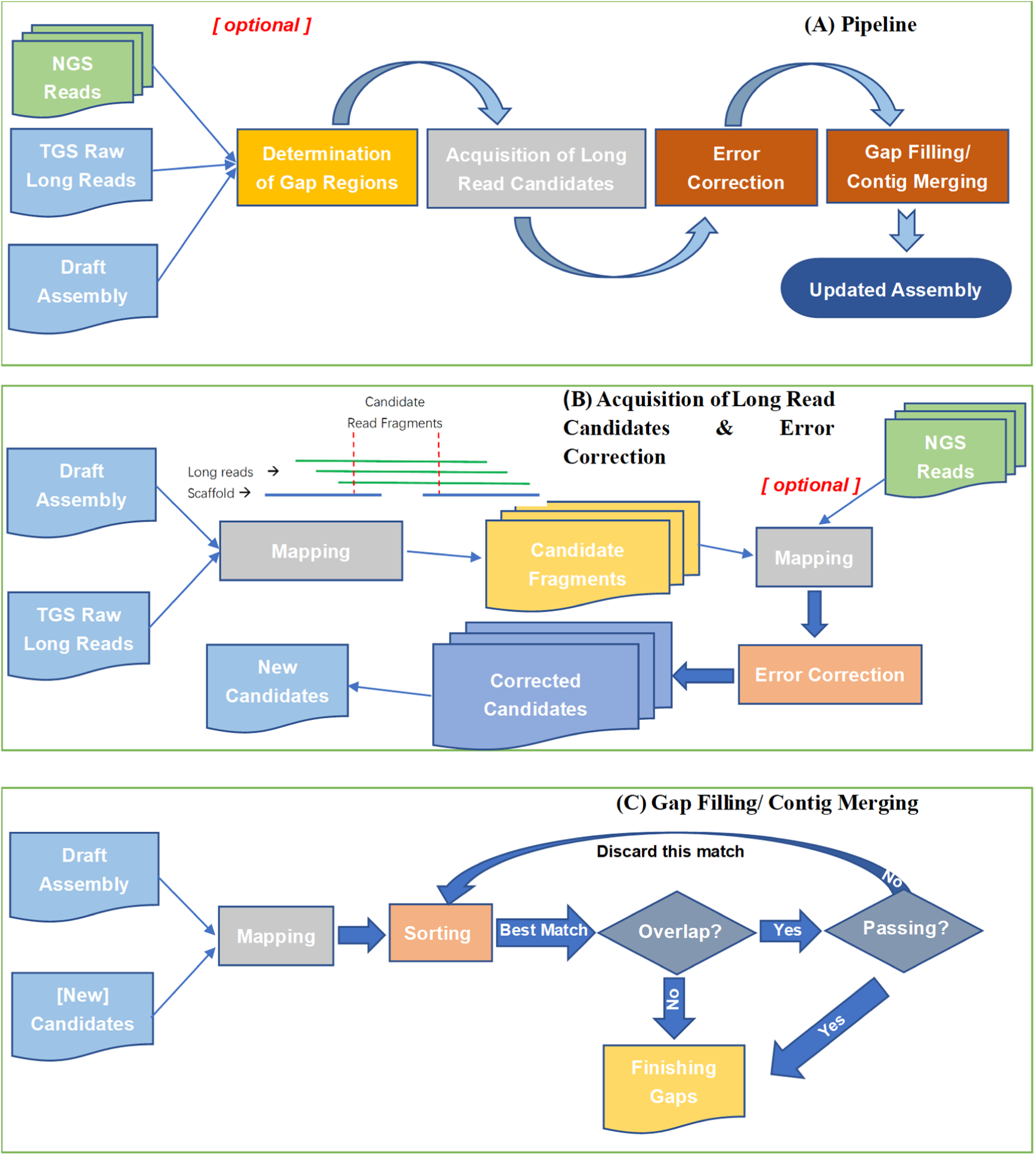
A schematic of TGS-GapCloser workflow. (A) A flow chart of overall algorithm, (B) A schematic describing the classification of gap regions, determination of candidate long read fragments, and error correction, (C) A detailed flow chart for gap filling or contig merging in a gap region used the most proper long-range information.

#### 2.2.1 Determination of gap regions

The input scaffolds were firstly split into parts called scaftigs from the observed N position, and each two neighboring scaftigs according to their positions in the same scaffold were considered as a gap region waiting to be filled. TGS-GapCloser defaults the high the quality of inputs, including base-level accuracy, order and orientation of scaftigs, but not the estimated gap size. That is because the estimations based on the long-range information provided by SLR, Hi-C, or BioNano cannot reach a resolution below ∼10kb, leading to a high probability of faults especially for short gaps.

#### 2.2.2 Acquisition of long read candidates

We used minimap2[36] to align the long reads against each gap region to obtain corresponding candidate fragments with the preset option -x ava-ont. The candidate for a specific gap is defined as the segment truncated from the aligned long reads in the area between two contigs along with 2kb flanking wings on both contigs. Each long read might provide several candidate sequences dependent on its spanning length and base-calling accuracy, but was limited to give at most one candidate for the same gap region to overcome the redundant alignments due to the algorithm’s nature of the aligner and high error rate of long reads.

The alignment quality determined the efficiency and accuracy of gap closure. Thus, all alignments were filtered based on the aligned length and identity ratio. For each gap, up to ten sub-long reads with the highest scores were chosen as candidates for error correction, avoiding multi-alignments within the same area and dramatically suppressing the data amount for further analysis. The quality score (QS) is given by

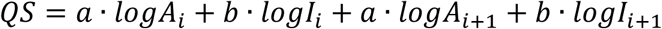

where letter *A* refers to the alignment length, letter *I* refers to the identity ratio for *i*th, *i*+*1*th contigs respectively in preliminary scaffolds; a and b are two arbitrary coefficients to distinguish A and I’s weights on the score, and have been tuned to 1:6 for ONT dataset as default. To further reduce the complexity and save the computational resources, the overlapped candidates in the same long read were clipped and merged.

#### 2.2.3 Error correction

The merged candidate sequences were error corrected using Pilon[37] to enhance both the base-level accuracy and the precision of alignments. Pilon tried to fix individual base errors, small indels and local misassemblies with short but accurate NGS reads. The short reads were aligned to the candidates also using minimap2 but conducted with the option -k14 -w5 -n2 -m20 -s 40 --sr --frag yes tuned for short sequences. Although Pilon was originally designed for assembly polishing and has a requirement of 50x or deeper genome coverage of short reads, the sequences in the waiting list were split into several groups and each group was then error corrected with the same NGS dataset to guarantee the coverage.

#### 2.2.4 Gap filling/contig merging

The error correction would benefit not only the single-base accuracy but also the final choice of the candidates to fill the gaps on the basis of the hypothesis that candidates with higher-quality alignments could be mapped to a more precise position in the reference after error correction, while those with lower-quality alignments became to fail to be mapped. The error-corrected candidates were again split and aligned against their corresponding gaps, and finally the one with the highest QS would be employed to fill the gap. We discarded the flanking wings of candidates but used bases from contigs as many as possible considering the relative accuracy.

If the best candidate gave a negative filling information instead, then the gap region would collapse to a single contig based on the overlapping relation. A portion of contigs have overlaps with others because of incorrect path in assembly graph or too aggressive contig extension strategy. But most scaffolders fail to deal with the overlapping relations, leaving a number of Ns to represent the uncertainty, and so do the majority of gap closers. However, a single long read spanning the gap has the ability to solve the overlapping if two contigs are mapped to the correct positions. We took more care of “negative” gaps given by the best corrected candidates with extra strict criteria because most base-calling errors in TGS long reads, including indels and homopoly-meric repeats tend to cause untruthful overlapping. Gaps without any corresponding candidate would fail to be closed.

### 2.3 Implementation

TGS-GapCloser was coded in C++ programing language. It applies minimap2 to align long reads against gaps or short reads against candidates, and Pilon (a Java package) to, which requires Java runtime 1.7 or later. The acceleration and enhanced mapping quality of the algorithm partially originate from the aligner, as minimap2 showed great improvement in speed and overall higher mapping accuracy for error-prone long reads, than others used in similar tools, such as BLASR[38] especially for unique and repetitive hits[36]. The algorithm automatically determined gap areas and tried to find the best matched long read fragments to fill gaps or merge adjacent contigs based on the alignments. The details in each step were individually recorded, including gap determination, mapping, long read extraction, error correction, and gap filling or merging. The final output was reported in FASTA format, along with log files describing the detailed sequence insertion or merging information for each gap to trace all the improvements. TGS-GapCloser is available via GitHub at https://github.com/BGI-Qingdao/TGSGapFiller.

### 2.4 Gap closing with other tools

We compared the performance with two of the most popular gap-closing tools, PBJelly (version PBSuite_15.8.24) and FGAP (version 1.8.1) using the same human Chr19 dataset. For PBJelly, gap closure was performed with raw ONT reads and the default options, while FGAP was conducted with the default options and overlap detection option off and on.

## 3 Results

### 3.1 Gap closure in human genome using different assemblies and long read datasets

Three genome assemblies and two long read datasets were used to assess the utility of TGS-GapCloser in gap closing or contig merging in preliminary scaffolds for *Homo sapiens* (NA12878). Using the same dataset, the whole genome was assembled by: (1) contigs by MaSuRCA[20] + scaffolds by SLR-superscaffolder[31]., (2) contigs by Mercedes+ scaffolds by SLR-superscaffolder, and (3) contigs and scaffolds by Supernova[33] to take full use of barcoded long-range information. Although MaSuRCA itself can assemble both contigs and scaffolds, the lack of stLFR information used in the assembler results in short scaffolds. It is necessary to employ SLR-superscaffolder to obtain comparable scaffold NG50 against Supernova. To declare the efficient usage of long reads, we randomly extracted ∼10 coverage from rel3[32] dataset with claimed 84.06% of median read identity[32]. The gap regions in the draft assemblies have been investigated to be fully covered by at least one long reads. Figure 2 describes the improvements after gap closure with our method. The contig NG 50 increased from 13.2kb to 593.4kb for assembly (1), 15.3kb to 660.7kb for assembly (2), and 109.6kb to 1193.4kb for assembly (3); while corresponding contig NGA50 grew from 13.1kb to 402.0kb, 15.2kb to 405.7kb, and 105.5kb to 724.9kb for three assemblies. Note that our current algorithm did not split or merge draft scaffolds to remain the existing long-range information. 91.8% of total 191,189, 94.8% of total 129,408, and 82.2% of total 42,359 gaps were successfully finished by TGS-GapCloser, respectively. After gap filling, genome fraction against the reference was improved by 1.4%, 3.1% and 0.4% for different inputs. The large-scale (>1kb) misassemblies were only increased by 17.4% and 9.7% in assembly (2) and (3), and even decreased by 10.7% in assembly (1) due to the more precise mapping position of scaffolds/contigs against the reference induced by the filling sequences. In spite of error correction, the local misassemblies (<1kb) still present an increment of 1.2-fold’, 7.4-fold, and 1.1-fold dependent on the single-base accuracy of filled sequences. Benchmarking Universal Single-Copy Orthologs (BUSCO)[39] (version 3.0.2) analysis indicated the possible enhancements for further analysis such as gene annotation after gap filling. The genome was queried against the vertebrata_odb9 database. It revealed that 90.5% [S: 89.3%, D:1.2%], 89.7% [S: 88.4%, D:1.3%], and 94.1% [S: 92.4%, D:1.7%] of the expected vertebrate genes were complete after gap filling, improved from the original 86.2% [S: 84.8%, D:1.4%], 76.6% [S: 75.6%, D:1.0%], and 90.7% [S: 89.1%, D:1.6%], respectively.

**Figure 2.**
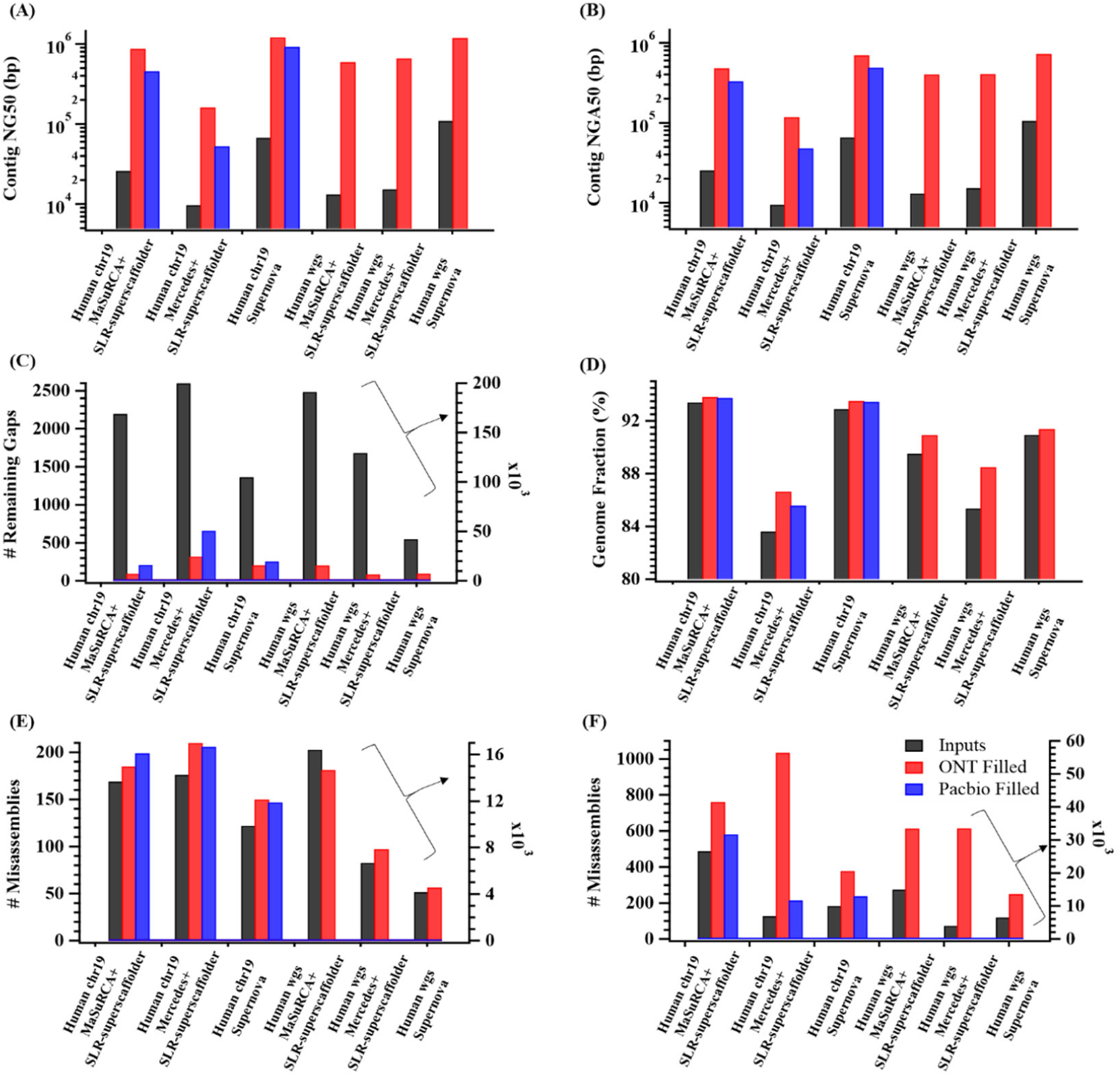
Gap filling improvements and effects on initial draft assembly produced by TGS-GapCloser for different inputs: (A) contig NG50, (B) contig NGA50, (C) number of remaining gaps, (D) genome fraction, (E) misassemblies and (F) local misassemblies estimated by QUAST[40].

The Chr19 datasets were extracted from the whole genome and assembled following the same steps for convenience. Using 28.6 mapped coverage of rel3 reads, the gap number in three preliminary scaffolds dropped to 97, 322, and 207, thus reducing the unknown regions by 95.6%, 87.6% and 84.8% for assembly (1), (2), and (3), respectively. The contig NG50 and NGA50 were also improved by 22.8 and 14.0 times on average. The increased genome coverage indicated that updated gap regions were mapped to reference’s new area. The increasing ratio of misassemblies and local misassemblies were consistent with the whole genome results, 17.2% and 2.9-fold. The gap length distribution of filled gaps is consistent with that of draft assemblies (Figure S1).

In addition, we tested the Pacbio dataset for Chr19 with 21.2 mapped coverage, with the acknowledged average read accuracy of 85%[21]. Applied to three assemblies with same parameters, TGS-GapCloser decreased the gap number by 90.4%, 74.7% and 81.1%, respectively. As a result, the contig NG50 and NGA50 were enhanced to 12.3- and 8.5-fold on average. After filling, the ratio of induced misassemblies and local misassemblies against inputs were 18.4% and 39.9%. The overall results were worse than those using ONT reads, for which shorter read length and lower read coverage might be responsible.

### 3.2 Gap closure in ultra large genome of ginkgo

Gingko is a best-known living fossil that has remained its form and structure over 270 million years, taking a unique position in the evolutionary tree of life[41]. We applied TGS-GapCloser to improve the chromosomal-level assembly of the *Ginkgo biloba*[34] by incorporating 11.9 coverage of error-corrected PacBio data. The updated assembly had been assigned to13 chromosomes of 9,570,195,624 bp, with 613,821 gaps in total. In this case, the long reads were pre-corrected by Canu. After gap filling which only consumed ∼541 CPU hours, 71.6% of the gaps were closed, and thus the total contig size increased by 411,608,879 bp, 4.3% of total. The contig N50 was also enhanced from 57.1kb to 364.8kb. Note that most tools have been only used for several bacterial and fungal genomes or small eukaryotes (<1Gb) previously[22, 25, 27], and it is doubt that it could be applied to this ultra large genome using reasonable computing resources.

### 3.3 Validation of gap-closing sequences

As a sanity check, we generated ideally filled sequences for all gaps using the reference of Chr19 and compared the theoretically filling results to gap sequences created by TGS-GapCloser. Table 1 lists the complete statistics for the evaluation of TGS-GapCloser’s improvements. By comparing actual filling sequences to the reference validated sequences, the PPV ranged from 99.1% for MaSuRCA+SLR-superscaffolder+ONT combinations to 77.3% for Supernova+ONT, and sensitivity from 87.4% to 55.2%, respectively.

**Table 1.** Gap Filling accuracy statistics for TGS-GapCloser. Sensitivity is defined as the ratio of the number of actually filled gaps that the reference also successively fills to the total number of gaps that the reference can fill. PPV is defined as the ratio of the number of actually filled gaps that can be uniquely mapped to the reference-filled gaps to the total number of filled gaps. Note that short sequences might introduce multi-alignments or mis-alignments, thus sequences shorter than 100bp were filtered out for this purpose. QV was evaluated in Phred format for all sequences instead. All datasets were run with 16 threads.

| Long Read Choosing Accuracy |
| --- |

| Input data | # of closed gaps | # of closed gaps in theory | PPV | Sensitivity | Runtime | Peak memory |
| --- | --- | --- | --- | --- | --- | --- |
| MaSuRCA+SLR-superscaffolder+ONT | 942 | 971 | 98.3% | 87.4% | 2h04min | 8.22 GB |
| Mercedes+SLR-superscaffolder+ONT | 1,335 | 1,565 | 99.1% | 79.3% | 1h01min | 7.41 GB |
| Supernova+ONT | 858 | 895 | 77.3% | 55.2% | 37min | 6.78 GB |

| Single Base Level Accuracy |  |  |  |  |  |
| --- | --- | --- | --- | --- | --- |
| Input data | # of closed bases (bp) | # of closed bases in theory (bp) | Input contig QV | Output contig QV | Filled QV |
| MaSuRCA+SLR-superscaffolder+ONT | 3,429,789 | 5,313,738 | 60.00 | 38.83 | 16.50 |
| Mercedes+SLR-superscaffolder+ONT | 8,015,998 | 14,861,676 | 42.84 | 30.20 | 15.55 |
| Supernova+ONT | 1,286,976 | 1,826,150 | 52.22 | 39.55 | 16.70 |

In terms of single-base level accuracy, the consensus quality value (QV) with the method in [32] was decreased by the inserted sequences, contig QV down from 51.7 to 36.2, with error-corrected filled QV of 16.3 on average. The final result with comparably lower single-base quality, however, was still better than the ONT final assembly QV, 21.5, assembled by Canu[35], error correction by nanopolish[42] and polishing by Pilon.

### 3.4 Comparison with other tools

We did not compare TGS-GapCloser to NGS gap-closing tools because the utilization of long/medium-range information provided by long reads spans repetitive or complicated regions that kmer-based extension cannot reach and congenitally creates better results as revealed in previous studies[22, 25]. In this work, we chose PBJelly and FGAP software in the gap closure analysis and applied to the Chr19 Mercedes+SLR-superscaffolder assembly with ONT dataset to compare gap filling performance. Other subsequent tools did not display obvious improvements in efficiency and accuracy of gap filling[27].

The evaluation of outputs showed that the gap-closing efficiency of TGS-GapCloser was considerably higher than that of other tools, leaving only 322 gaps compared to 1,730 for PBJelly default, 1,016 gaps for FGAP default and 782 for FGAP with overlap option on (Table 2), thus enhancing the contig NG50 and NGA50 from 11.4kb to 165.2kb and 9.3kb to 117.6kb, respectively, 3.4-6.1-fold than PBJelly and FGAP. The induced misassemblies were less than that of PBJelly and FGAP default but comparable with FGAP (overlap on), whereas FGAP (overlap on) showed the least local misassemblies. Considering the common existence of overlapped adjacent contigs in input scaffolds, the overall result of FGAP with overlap option on was better than that of default settings. The number of mismatches indels per 100kb varied, but larger than that of draft scaffolds, especially for the indels because of the nature of ONT sequencing platform. Note that PBJelly might elongate scaffolds by local reassembly but required sufficient data coverage.

**Table 2.**
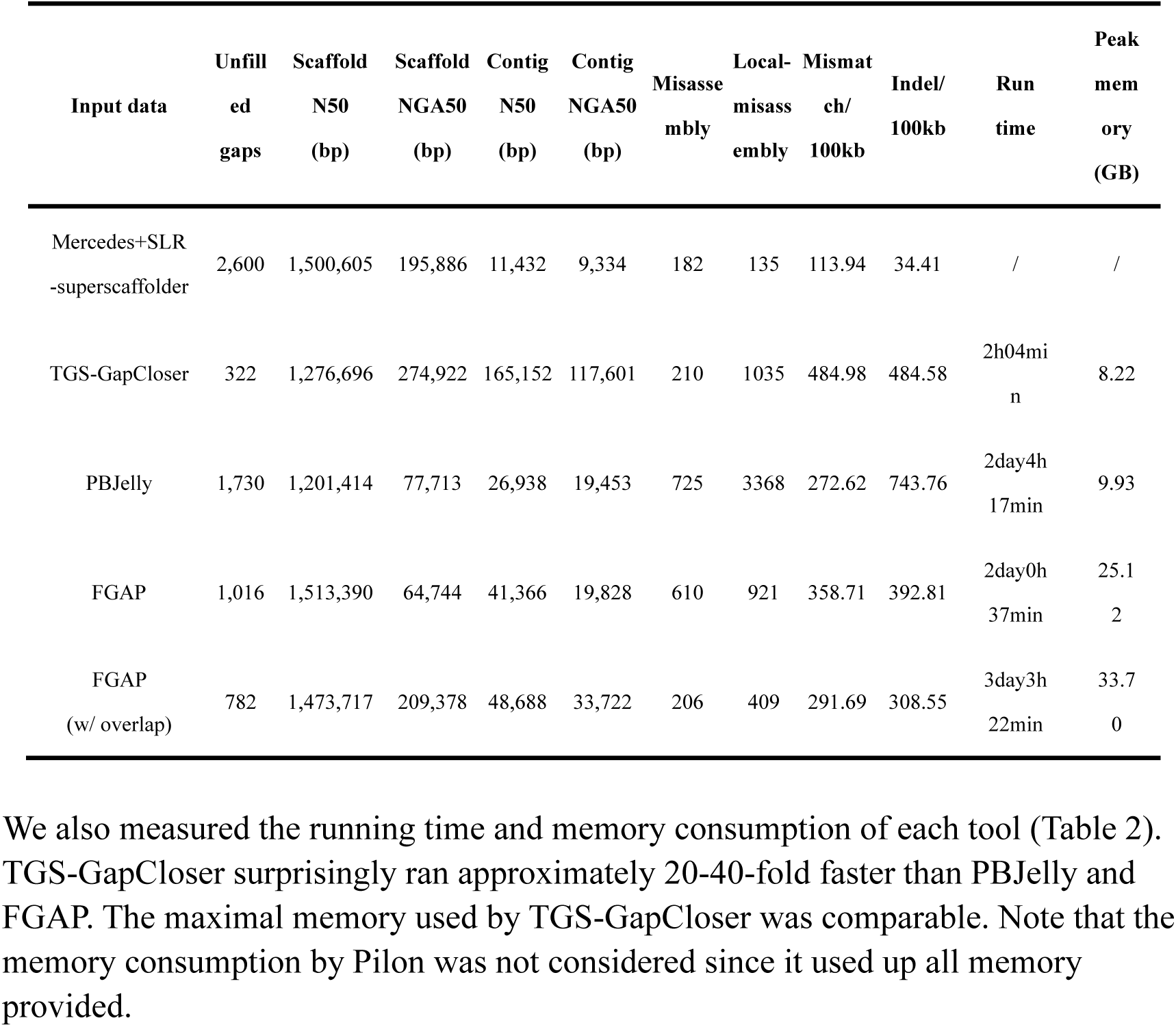
Gap filling statistics for TGS-GapCloser, PBJelly and FGAP. All datasets were run with 16 threads.

## 4 Discussion

### 4.1 Computational resource consumption

We have presented a new tool for updating a draft genome assembly based on currently available long reads fast and accurately. This method is widely applicable to many genome projects for thousands of research groups, benefiting from its flexibility of using different sequencing technologies or different assemblies. TGS-GapCloser requires only low coverage of expensive long reads without pre-error correction, making this approach more costly effective and suitable for small budgets. With respect to the human whole genome, it consumed 4,498 CPU hours in total and the peak memory was 58.1 GB excluding Pilon on average, correcting only relatively short fragments in gap regions. Even Pilon ran successfully with 1TB memory provided for a 3.2 Gb genome, although it was recommended 3.2 TB at least[37]. The gap-closing algorithm is more costly effective compared with the *de novo* assembly of 30× long reads with Canu, which requires ∼40K CPU hours for ONT and ∼62K CPU hours for Pacbio[32]. The speed and memory usage are further improved without error correction. It took only ∼541 CPU hours for ginkgo’s large genome using pre-error-corrected Pacbio reads.

### 4.2 Future direction

A number of improvements for future versions of TGS-GapCloser have been put on the agenda. The accuracy of inserted sequences largely depends on the performance of the aligner. Minimap2 performs well in most cases, however, gets worse when handing with a small portion of overlapping relations for long sequences and short reads against long sequences although it has been tuned somewhat. We hope that the problem will be solved by applying other aligners or sufficient parameter tuning. In addition, the computational consumption by Pilon is still considerable although the algorithm has tried to reduce the input data size as much as possible. It is convenient to replace it with other error-correction tools when available. Longer reads with higher quality are promised by ONT and Pacbio, which help us get rid of this annoying step. Last but not least, we trusted the input scaffolds as well as the orientation and order of contigs in each scaffold to retain the existing assembly information, but neglected the assembly errors in reality. We are planning to use the long/medium-range information provided by TGS reads to correct the improper relation of contigs in the same scaffold and link different scaffolds if there is overlapping.

## Acknowledgements

The authors are grateful for the advice from Hongmei Zhu and many other BGI-Shenzhen employees in the development of TGS-GapCloser.

