## Supplementary_Information for "TGS-GapCloser: fast and accurately passing through the Bermuda in large genome using error-prone third-generation long reads"

### 1. Genome assemblies and TGS datasets

Three datasets of two large genomes were used to examine the gap-closing results by TGS GapCloser. We sequenced *Homo sapiens* (NA12878) using MGIEasy single tube-Long Fragment Reads (stLFR) Library Prep Kit and BGISEQ-500 with the data size of 660 Gb, and reads mapped to the Chromosome 19 (Chr19) were also extracted for further analysis. NGS short reads were assembled using de Bruijn graph-based assemblers, MaSuRCA(*1*) and Mercedes (unpublished) to obtain short but highly accurate contigs for each dataset, and the SLR long-range and paired-end information provided by stLFR technique was exploited to do further scaffolding by SLR-superscaffolder(*2*). Supernova(*3*) was originally designed to assemble 10X Genomics data, but could be applied to stLFR format reads to obtain draft scaffolds. To test the generalization of TGS Gapcloser, we utilized both Pacbio RSII (SRR3197748) downloaded from GIAB and ONT MinION (rel3)(*4*) long reads to close the gaps. The input genome assembly of *Ginkgo biloba* female (estimated 10.61 Gb) was obtained from (*5*), which assembled using SOAPdenovo2(*6*) and updated using Hi-C data. The Pacbio reads for ginkgo were sequenced by Pacbio Sequel, with chemistry of Sequel Sequencing Kit 3.0 Bundle (4 rxn). The total data amount was 256 Gb with the average read length of 38,623 bp. Error-correction by Canu(*7*) reduced data to 126 Gb, with the average read length of 10,722 bp.

### 2. Validation of gap-closing results

We classified the elevations of the gap-closing accuracy in two levels: the long read choosing accuracy and the single-base accuracy. The former was determined by whether the algorithm captured the best matched error-prone long reads to the corresponding gap region, and affected the large/medium assembly information, such as chromosomal variations. The quality of error correction decided the latter accuracy, and affected the small single-nucleotide polymorphism or insertion/deletion calling in short range.

QUAST(*8*) (version 5.0.2) generated basic assembly statistics, as well as misassemblies (>1kb), local misassemblies (<1kb), mismatches and indels for errors at different levels when a reference genome was provided. To further evaluate the efficiency and accuracy, we aligned the reference against the assembly to get the theoretically filled gap sequences with QUAST intermediate files, and compared them to sequences filled by TGS Gapcloser with minimap2 (preset -x map-ont). The sensitivity was defined to the ratio of the filled gap number to the total gap number in input scaffolds which had corresponding sequences in the reference, and the PPV was defined to the ratio of the number of gaps for which the filled and reference sequences were matched to that of total filled gaps with corresponding reference to judge. The single base level accuracy was assessed by mapping the NGS reads to sequences to evaluate the QV with the method in (*4*). The QVs were expressed in Phred format.

**3. Long read coverage effect**

It is worthwhile evaluating the effect of long read coverage on gap filling results. We extracted 1×, 5×, 10×, 20× and original 29× mapped ONT rel3 reads for Chr19, and individually ran the gap closure with them using the same default options. As shown in the Figure 3 (A), the number of filled gap and total filled bases grow with the increasing coverage, and saturates at 10× to the level of theoretically filled gap number and sequences. Surprisingly, the total time usage does not change much as the coverage increases, while the peak memory presents an approximately linear growth in Figure 3 (B). With more coverage of long reads, Figure 3 (C) displays that the sensitivity increases from 22.1% to 87.4% while the PPV remains similar. In terms of single-base level accuracy in Figure 3 (D), the QV of inserted sequences drops as more gaps are filled, but has ignorable effect on that of the output contigs. The result indicates that TGS GapCloser enables a considerable number of gaps filled using low coverage of long reads.

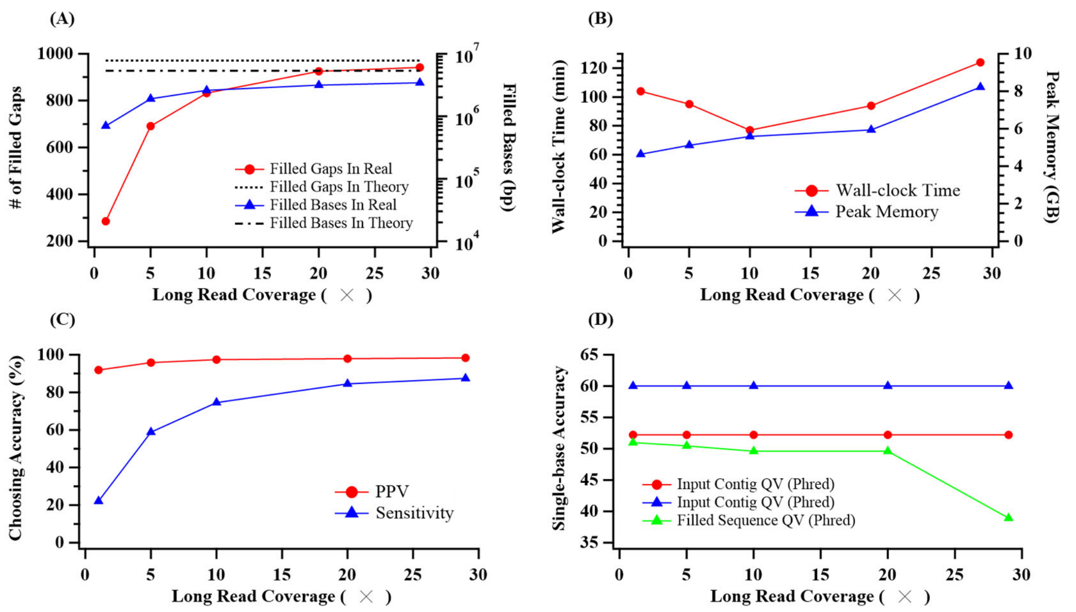


**Figure S1. Long read coverage effect on gap filling:** (A) number of filled gaps and bases, (B) wall-clock time and peak memory, (C) choosing accuracy for long reads, and (D) single-base level accuracy. All datasets were run with 16 threads.

### 4. Filled Gap length distribution

The gaps in different draft scaffolds exhibit different length distribution as shown in Figure S2, however the majority is located in either positive or negative short length range. The length distribution of filled gaps behaves highly similarly to that of corresponding inputs for different scenarios. But the negative short gaps are slightly overfilled by TGS GapCloser, possibly because of the bad alignment quality for neighboring contigs with small overlapping portion found by the aligner.

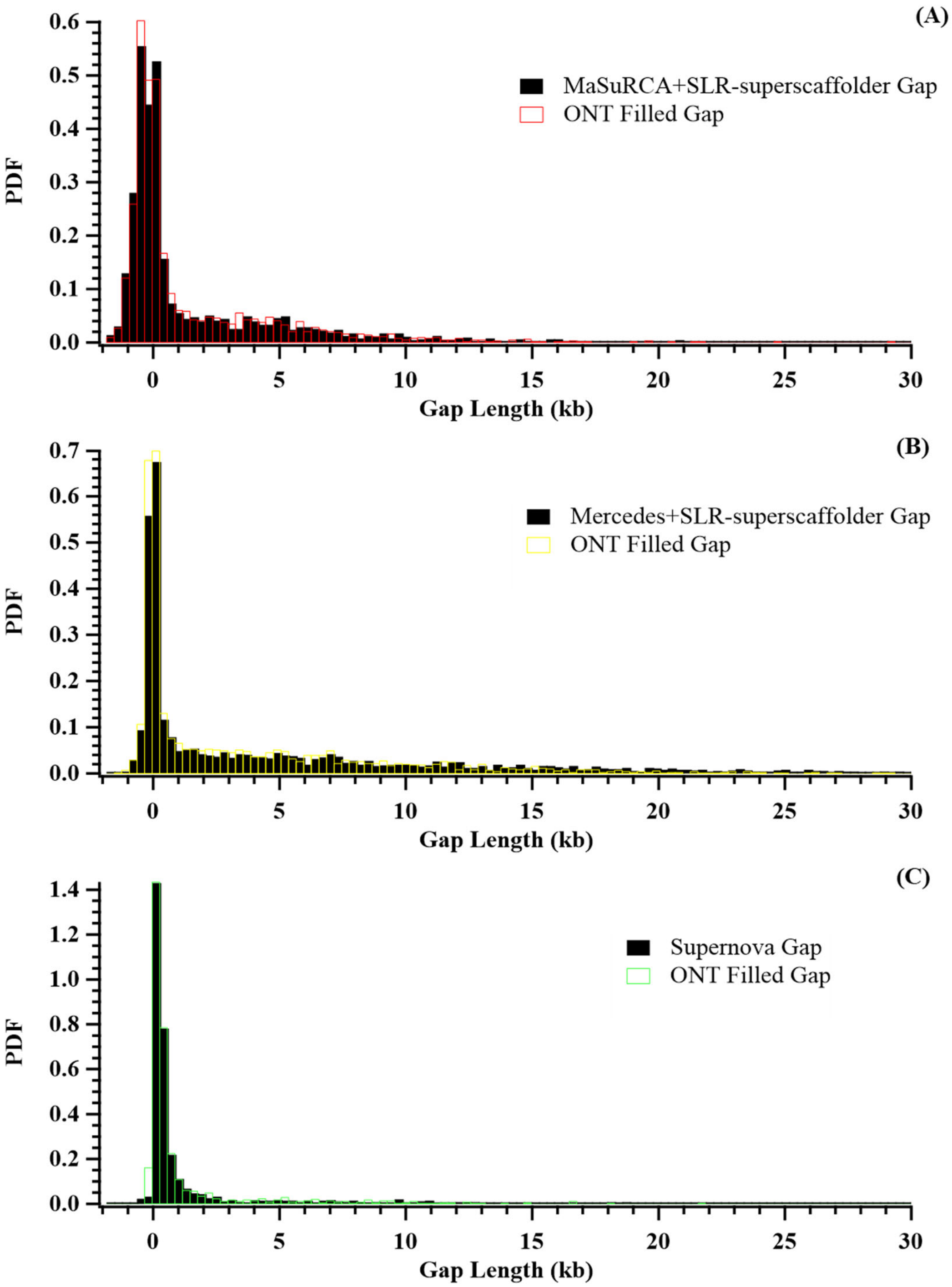


**Figure S2. Length distribution of gaps in initial scaffolds and filled by TGS GapCloser.**

**Table S1. Summary of the input contigs/scaffolds in this work**

| Organisms | Datasets | Total length | Genome Fraction | # of Scaffolds | # of gaps | Scaffold NG50/N50 (bp) | Scaffold NGA50 (bp) | Contig NG50/N50 (bp) | Contig NGA50 (bp) | # of misassemblies | # of local misassemblies | BUSCO |
|---|---|---|---|---|---|---|---|---|---|---|---|---|
| Human Chr19 | MaSuRCA+SLR-superscaffolder | 62,941,506 | 93.405% | 2,202 | 2,195 | 8,696,596 | 873,719 | 25,956 | 25,384 | 169 | 488 | / |
| | Mercedes+SLR-superscaffolder | 66,644,779 | 83.62% | 3,411 | 2,600 | 1,561,142 | 196,307 | 9,687 | 9,464 | 176 | 126 | / |
| | Supernova | 57,459,696 | 92.901% | 1,631 | 1,363 | 8,378,184 | 964,978 | 67,317 | 65,678 | 122 | 183 | / |
| *Homo sapiens* (NA12878) | MaSuRCA+SLR-superscaffolder | 3,345,341,888 | 89.514% | 142,241 | 191,189 | 17,657,864 | 380,495 | 13,216 | 13,101 | 16,426 | 15,012 | C:86.2%[S:84.8%,D:1.4%],F:8.4%,M:5.4% |
| | Mercedes+SLR-superscaffolder | 3,201,977,072 | 85.360% | 72,359 | 129,408 | 11,800,332 | 568,904 | 15,295 | 15,210 | 6,699 | 3,990 | C:76.6%[S:75.6%,D:1.0%],F:13.8%,M:9.6% |
| | Supernova | 2,855,133,582 | 90.939% | 40,738 | 42,359 | 26,284,488 | 1,837,192 | 109,644 | 105,516 | 4,172 | 6,509 | C:90.7%[S:89.1%,D:1.6%],F:4.5%,M:4.8% |
| *Ginkgo biloba* | SOAPdenovo2+HiC | 9,570,195,624 | / | 13 | 613,821 | 724,988,903 | / | 57,099 | / | / | / | |

**Table S2. Summary of the updated contigs/scaffolds in this work**

| Organisms | Input data | Contig NG50/N50 (bp) | Contig NGA50 (bp) | # of closed gaps | # of filled bases (bp) | Genome fraction | Misassemblies | Local Misassemblies | BUSCO |
|---|---|---|---|---|---|---|---|---|---|
| Human chr19 | MaSuRCA+SLR-superscaffolder+ONT | 870,665 | 479,099 | 2,098 | 2,759,626 | 93.802% | 185 | 762 | / |
| | Mercedes+SLR-superscaffolder+ONT | 161,606 | 117,601 | 2,278 | 7,750,957 | 86.640% | 210 | 1035 | / |
| | Supernova+ONT | 1,215,091 | 699,712 | 1,156 | 1,254,168 | 93.517% | 150 | 378 | / |
| | MaSuRCA+SLR-superscaffolder+Pacbio | 456,879 | 330,145 | 1,984 | 1,752,579 | 93.746% | 199 | 582 | / |
| | Mercedes+SLR-superscaffolder+Pacbio | 52,981 | 47,978 | 1,942 | 3,465,833 | 85.590% | 206 | 215 | / |
| | Supernova+Pacbio | 923,687 | 489,067 | 1,106 | 794,976 | 93.450% | 147 | 238 | / |
| *Homo sapiens* (NA12878) | MaSuRCA+SLR-superscaffolder+ONT | 593,445 | 401,962 | 175,437 | 272,947,305 | 90.922% | 14,667 | 33,477 | C:90.5%[S:89.3%,D:1.2%],F:5.2%,M:4.3% |
| | Mercedes+SLR-superscaffolder+ONT | 660,736 | 405,727 | 122,626 | 336,390,884 | 88.490% | 7,862 | 33,536 | C:89.7%[S:88.4%,D:1.3%],F:5.8%,M:4.5% |
| | Supernova+ONT | 1,193,354 | 724,876 | 34,831 | 48,546,731 | 91.372% | 4,576 | 13,636 | C:94.1%[S:92.4%,D:1.7%],F:2.7%,M:3.2% |
| *Ginkgo biloba* | SOAPdenovo2+Hic+Pacbio | 364,800 | / | 439,641 | 411,608,879 | / | / | / | |

Table S3. Genomics datasets source

| Organisms | Datatype | Total reads | Total bases | Avg. read length | Source |
|---|---|---|---|---|---|
| Human Chr19 | stLFR | 43,014,410 | 4,301 Mb | PE100 | Extracted from *H. sapiens* |
| | ONT | 250,811 | 1,673 Mb | 6,671 | Extracted from *H. sapiens* |
| | Pacbio | 222,699 | 1,243 Mb | 5,583 | Extracted from *H. sapiens* |
| | Reference | 1 | 58,617,616 bp | 58,617,616 | Extracted from *H. sapiens* |
| *Homo sapiens* (NA12878) | stLFR | 2,071,968,776 | 660 Gb | PE100 | ftp://ftp.cngb.org/pub/CNSA/CNP0000066/CNS0007594/CNX0005851/CNR0006062/ |
| | ONT | 14,183,584 | 91 Gb | 6,433 | https://github.com/nanopore-wgs-consortium/NA12878/blob/master/nanopore-human-genome/rel_3_4.md |
| | Pacbio | 13,099,385 | 75 Gb | 5,696 | ftp://ftp-trace.ncbi.nih.gov/giab/ftp/data/NA12878/NA12878_PacBio_MtSinai/sorted_final_merged.bam |
| | Reference | 455 | 3,209,286,105 bp | 7,053,376 | ftp://ftp.ncbi.nlm.nih.gov/genomes/all/GCF/000/001/405/GCF_000001405.39_GRCh38.p13/GCF_000001405.39_GRCh38.p13_genomic.fna.gz |
| *Ginkgo biloba* | NGS Dataset | 18,213,533,333 | 1,690 Gb | PE150/PE100/PE75 | https://www.ncbi.nlm.nih.gov/sra?linkname=bioproject_sra_all&from_uid=307642 |
| | Pacbio | 6,717,458 | 256 Gb | 38,623 | https://doi.org/10.5524/100613 |

**Table S4. Control parameters used for different datasets.**

| Software | Parameters |
| --- | --- |
| MaSuRCA | GRAPH_KMER_SIZE=63, cgwErrorRate=0.15 |
| Mercedes | default |
| SLR_superscaffolder | MST_BIN_SIZE=7000, HT_BIN_SIZE=3500, CLUSTER=0.1, PE_SEED_MIN=200 |
| Supernova | --nopreflight |
| TGS GapCloser | CHUNK_NUM=3, MinIDY=0.3, MinMatch=300 |
| PBJelly | --minMatch 8 --minPctIdentity 70 --bestn 1 --nCandidates 20 --noSplitSubreads |
| FGAP | -R 100000 -I 100000 –p 1 –z 0 –g 0 |
| FGAP (w/ overlap) | -R 100000 -I 100000 –p 1 –z 1 –g 1 |
| QUAST | -m1000 –s --fast --fragmented |
| BUSCO | -l vertebrata_odb9 −m geno −sp human |
